# SARS-CoV-2 Delta breakthrough infections in vaccinated patients

**DOI:** 10.1101/2022.04.12.488092

**Authors:** Jing Zou, Xuping Xie, Mingru Liu, Pei-Yong Shi, Ping Ren

**Affiliations:** Department of Biochemistry and Molecular Biology, University of Texas Medical Branch, Galveston TX; Department of Pathology, University of Texas Medical Branch, Galveston TX; Institute for Human Infection and Immunity, University of Texas Medical Branch, Galveston, TX; Sealy Institute for Drug Discovery, University of Texas Medical Branch, Galveston, TX; Institute for Translational Sciences, University of Texas Medical Branch, Galveston, TX; Sealy Institute for Vaccine Sciences, University of Texas Medical Branch, Galveston, TX; Sealy Center for Structural Biology & Molecular Biophysics, University of Texas Medical Branch, Galveston, TX

**Keywords:** COVID-19, SARS-CoV-2, breakthrough infection, antibody neutralization, vaccine, variants of concern, vaccine booster

## Abstract

The continuous emergence of SARS-CoV-2 variants with increased transmission and immune evasion has caused breakthrough infections in vaccinated population. It is important to determine the threshold of neutralizing antibody titers that permit breakthrough infections. Here we tested the neutralization titers of vaccinated patients who contracted Delta variant. All 75 patients with Delta breakthrough infections exhibited neutralization titers (NT_50_) of less than 70. Among the breakthrough patients, 76%, 18.7%, and 5.3% of them had the NT_50_ ranges of <20, 20-50, and 50-69, respectively. These clinical laboratory results have implications in vaccine strategy and public health policy.

## Main text

The COVID-19 pandemic is umpired by two dynamic factors: (i) the continuous emergence of SARS-CoV-2 variants with improved transmission and/or immune evasion and (ii) the waning immunity post vaccination and infection. This is exemplified by the two recent surges of Delta and Omicron variants, which caused many breakthrough infections in vaccinated individuals. Since antibody neutralization is a key contributor to vaccine protection, it is thus important to define the neutralization levels in patients with breakthrough infections. Such information is essential to guide vaccine strategy and policy. Here we characterized the antibody neutralization in vaccinated patients when they contracted Delta variant infections.

To determine the neutralization titers (NT_50_) in breakthrough patients when they got infected with Delta variant, we collected sera from 75 patients who were vaccinated and subsequently contracted breakthrough infections. **Table 1** summarizes the patient information and their NT_50_ values. All patients were vaccinated with 2 doses of Pfizer or Moderna vaccine or 1 dose of J&J vaccine. Breakthrough infections were confirmed by positive viral RNA tests. Although the genotypes of individual infecting viruses were not determined by sequencing, they were most likely Delta variant because all infections had occurred from late July to October, 2021 when Delta was 100% prevalent in our patient population based on the local SARS-CoV-2 surveillance system and about 98% prevalent in the USA (https://covid.cdc.gov/covid-data-tracker/#variant-proportions). All sera were taken 0-5 days before the positive viral nucleic acid tests. We determined the NT_50_ of each serum using a well-established mNeonGreen reporter USA/WA1/2020 SARS-CoV-2 [1]. This neutralization assay has been reliably used to support the BNT162b2 vaccine development [2]. The NT_50_ value was defined as the interpolated reciprocal of the dilution yielding a 50% reduction in mNeonGreen-positive cells. Each specimen was tested in duplicates and the geometric mean of the duplicate results is presented. The first serum dilution of the neutralization test was 1:20. The NT_50_ values of any specimens with no detectable neutralizing activities at 1:20 dilution were treated as 10 for plot and calculation purposes (**Table 1**). The overall results reveal three observations. *First*, all breakthrough patients had low NT_50_s of <70 (**FIG 1A**). About 76%, 18.7%, and 5.3% of the breakthrough patients exhibited the NT_50_ ranges of <20, 20-50, and 50-69, respectively (**FIG 1B**). The results suggest NT_50_ of 70 as a neutralizing threshold required to prevent Delta breakthrough infections. *Second*, senior people appeared to be more vulnerable to breakthrough infections. Approximately 16%, 25.3%, and 58.7% of the breakthrough cases were in the age groups of 20-40, 41-64, and 65-97, respectively (**FIG 1C**). However, the NT_50_ differences among the three age groups are not all statistically significant (**FIG 1D**). *Third*, almost 90% of the breakthrough patients had received the 2 doses Pfizer or Moderna vaccine or 1 dose J&J vaccine for more than 120 days (**FIG 1E**). However, this observation was not statistically correlated with the NT_50_ differences among different time frames post-vaccination (**FIG 1F**).

**Table 1.** Serum information and NT_50_values

| Serum ID | Age (years) | Gender (F/M) | Ethnicity | Serum collection time (days post positive *NAAT) | Vaccine type | #Days post last vaccination | <sup>^</sup>NT<sub>50</sub> |
| --- | --- | --- | --- | --- | --- | --- | --- |
| 1 | 94 | F | Black | 0 | Pfizer | 137 | <sup>&</sup>10 |
| 2 | 74 | F | White | 0 | Pfizer | 176 | 10 |
| 3 | 74 | F | White | -4 | Pfizer | 193 | 10 |
| 4 | 91 | F | White | 0 | Pfizer | 203 | 10 |
| 5 | 83 | F | White | 0 | Pfizer | 182 | 10 |
| 6 | 34 | M | White | 0 | Pfizer | 139 | 10 |
| 7 | 66 | M | Black | 0 | Pfizer | 178 | 10 |
| 8 | 60 | M | Hispanic | 0 | Moderna | 159 | 10 |
| 9 | 64 | F | Black | 0 | Pfizer | 124 | 10 |
| 10 | 76 | M | White | 0 | Pfizer | 19 | 10 |
| 11 | 42 | F | Black | -5 | Pfizer | 216 | 10 |
| 12 | 53 | F | White | -4 | Pfizer | 149 | 10 |
| 13 | 79 | F | Black | 0 | Pfizer | 149 | 10 |
| 14 | 50 | M | Black | -1 | Pfizer | 101 | 10 |
| 15 | 85 | M | White | 0 | Pfizer | 187 | 10 |
| 16 | 72 | M | Black | 0 | Pfizer | 186 | 10 |
| 17 | 72 | F | Black | 0 | Pfizer | 166 | 10 |
| 18 | 88 | M | White | 0 | Moderna | 207 | 10 |
| 19 | 34 | F | White | -3 | Pfizer | 128 | 10 |
| 20 | 56 | F | White | 0 | Pfizer | 224 | 10 |
| 21 | 44 | F | White | 0 | Pfizer | 225 | 10 |
| 22 | 77 | F | White | 0 | <sup>†</sup>J&J | 179 | 10 |
| 23 | 97 | F | White | 0 | Pfizer | 205 | 10 |
| 24 | 54 | F | Black | -1 | Pfizer | 120 | 10 |
| 25 | 80 | F | Hispanic | 0 | Pfizer | 196 | 10 |
| 26 | 84 | M | White | 0 | Pfizer | 198 | 10 |
| 27 | 73 | F | White | 0 | Pfizer | 190 | 10 |
| 28 | 68 | M | White | -3 | Pfizer | 231 | 10 |
| 29 | 69 | M | White | 0 | Pfizer | 195 | 10 |
| 30 | 30 | M | White | 0 | Moderna | 156 | 10 |
| 31 | 52 | F | White | 0 | Pfizer | 98 | 10 |
| 32 | 69 | F | Black | 0 | J&J | 188 | 10 |
| 33 | 75 | F | White | 0 | Pfizer | 165 | 10 |
| 34 | 73 | M | White | -3 | Pfizer | 234 | 10 |
| 35 | 73 | F | White | 0 | Pfizer | 179 | 10 |
| 36 | 62 | F | Hispanic | 0 | Pfizer | 225 | 10 |
| 37 | 68 | F | Black | 0 | J&J | 212 | 10 |
| 38 | 59 | M | Black | 0 | Pfizer | 149 | 10 |
| 39 | 80 | M | Hispanic | 0 | J&J | 170 | 10 |
| 40 | 77 | F | Asian | 0 | Pfizer | 236 | 10 |
| 41 | 35 | M | White | 0 | Pfizer | 138 | 10 |
| 42 | 72 | M | White | 0 | Pfizer | 199 | 10 |
| 43 | 73 | M | White | -5 | Pfizer | 219 | 10 |
| 44 | 87 | M | White | 0 | Pfizer | 223 | 10 |
| 45 | 84 | F | Black | 0 | J&J | 211 | 10 |
| 46 | 85 | F | White | 0 | Pfizer | 241 | 10 |
| 47 | 78 | F | Hispanic | 0 | Pfizer | 236 | 10 |
| 48 | 89 | M | White | 0 | Pfizer | 162 | 10 |
| 49 | 56 | M | Hispanic | 0 | Moderna | 154 | 12 |
| 50 | 77 | M | White | 0 | Moderna | 238 | 14 |
| 51 | 77 | M | Black | 0 | J&J | 165 | 14 |
| 52 | 96 | F | Black | 0 | J&J | 218 | 15 |
| 53 | 66 | M | White | 0 | Pfizer | 178 | 15 |
| 54 | 85 | F | Black | 0 | Pfizer | 161 | 16 |
| 55 | 64 | M | White | 0 | Pfizer | 156 | 19 |
| 56 | 65 | M | Black | 0 | Pfizer | 113 | 19 |
| 57 | 30 | M | White | -3 | Pfizer | 223 | 19 |
| 58 | 85 | F | White | 0 | Pfizer | 194 | 20 |
| 59 | 62 | M | White | -1 | Pfizer | 175 | 20 |
| 60 | 60 | F | White | 0 | Pfizer | 41 | 21 |
| 61 | 46 | F | Black | 0 | Pfizer | 173 | 21 |
| 62 | 16 | F | White | -1 | Pfizer | 129 | 21 |
| 63 | 20 | F | White | 0 | Pfizer | 119 | 22 |
| 64 | 41 | F | White | -4 | Pfizer | 190 | 26 |
| 65 | 34 | F | Hispanic | 0 | Moderna | 163 | 27 |
| 66 | 86 | F | White | 0 | Moderna | 187 | 31 |
| 67 | 59 | M | White | 0 | Pfizer | 142 | 33 |
| 68 | 61 | M | Native Hawaiian | -3 | Pfizer | 175 | 35 |
| 69 | 80 | F | White | 0 | Pfizer | 193 | 41 |
| 70 | 38 | M | Hispanic | 0 | Pfizer | 104 | 41 |
| 71 | 27 | M | Hispanic | 0 | Pfizer | 145 | 47 |
| 72 | 89 | M | White | 0 | Pfizer | 195 | 51 |
| 73 | 39 | F | White | -3 | Pfizer | 181 | 52 |
| 74 | 91 | F | White | 0 | Pfizer | 166 | 65 |
| 75 | 38 | F | Hispanic | 0 | Pfizer | 69 | 69 |

\*Nucleic acid amplification test (NAAT)
#Days after dose 2 of Pfizer and Moderna vaccine or days after dose 1 of J&J vaccine
^Individual NT<sub>50</sub> value is the geometric mean of duplicate neutralization test results.

**FIG 1.**
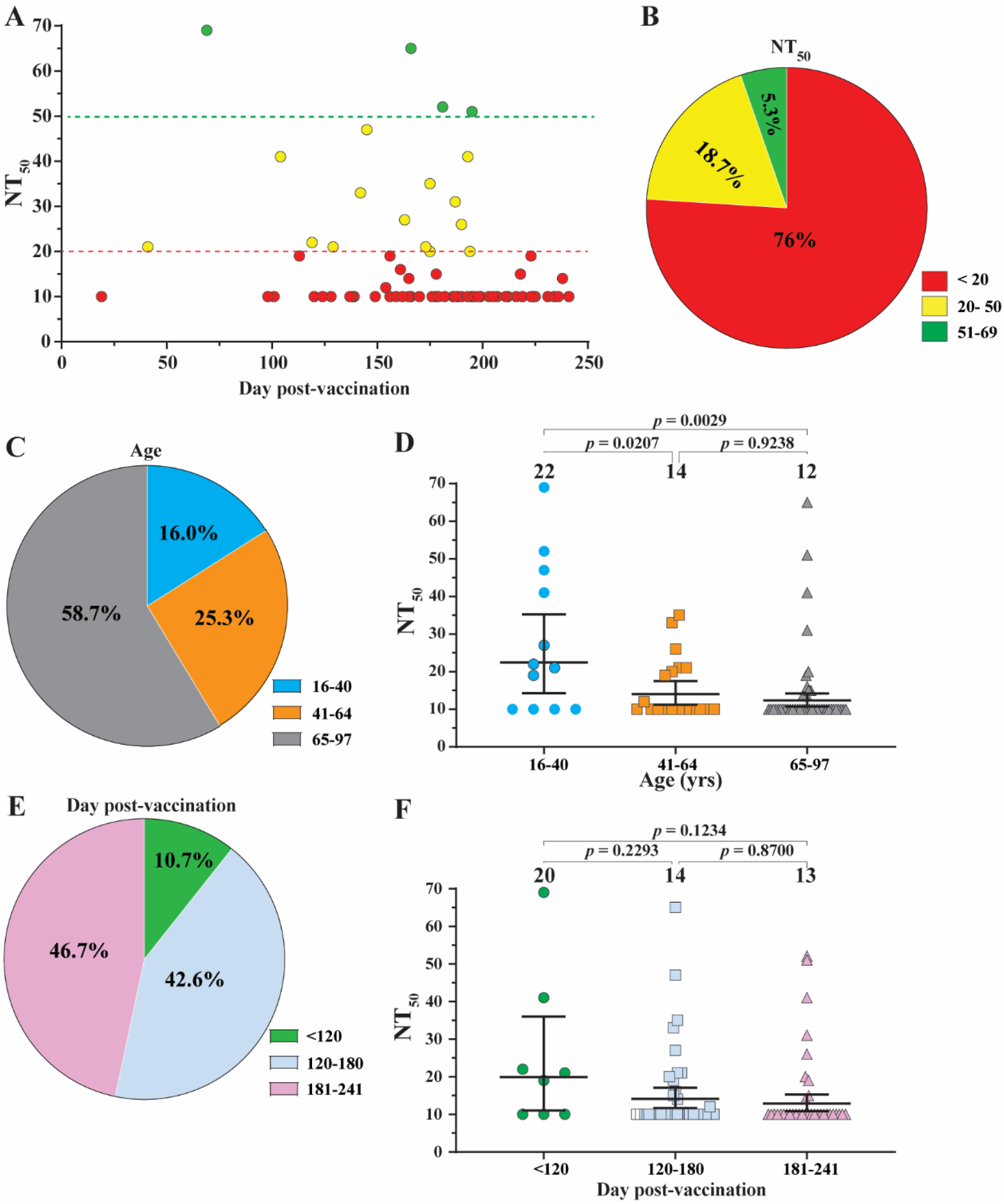
Analysis of Delta breakthrough infections in vaccinated patients. A panel of 75 sera, collected from Delta breakthrough patients, were measured for antibody neutralization titers (NT_50_) against USA-WA1/202. All patients were immunized with 2 does of Pfizer or Moderna vaccine or 1 dose of J&J vaccine. All sera were taken 0-5 days before viral RNA-positive test results. All patient information and NT_50_ values are detailed in **Table 1. A**. Plot of NT_50_ values versus days after dose 2 of Pfizer or Moderna vaccine or after 1 dose of J&J vaccine. Each data point represents the geometric mean of NT_50_ for one serum tested in duplicate assays. Different colors represent different NT_50_ ranges. Samples with no detectable neutralizing activities were plotted as 10 for calculation purpose. **B**. Pie presentation of different NT_50_ ranges. **C**. Age distributions. **D**. Plot of NT_50_ of different age groups. **E**. Distribution of breakthrough percentages versus days post-vaccination. **F**. Plot of NT_50_ versus days post-vaccination. In **D** and **F**, the bar heights and the numbers above indicate geometric mean titers. The whiskers indicate 95% confidence intervals. Statistical analysis was performed with the use of the one-way ANOVA with Tukey’s correction for multiple comparisons test.

One limitation of this study is that the NT_50_ values were measured against the original USA-WA1/2020 SARS-CoV-2 (a strain isolated in late January 2020), not directly against the delta variant. Previous studies showed that BNT162b2-vaccinated sera (collected at 1 month after dose 2) neutralized Delta variant at an efficiency that was 31-69% lower than the USA-WA1/2020 [3]. Compared with Delta, the newly emerged Omicron is significantly less susceptible to neutralization by vaccinated or non-Omicron infected human sera [4, 5]. The reduced neutralization susceptibility, combined with the increased transmissibility of Omicron, may have accounted for the high breakthrough infections observed in the current Omicron surge. These laboratory investigations, together with the real-world vaccine effectiveness, will provide guidance for future vaccine strategy and public health policy.

## Acknowledgements

We thank Michael L O’Rourke from the Information System Department at UTMB for assisting with electronic medical record systems.

